# Developing Hybrid Systems to Address O_2_ Uncoupling in Multi-Component Rieske Oxygenases

**DOI:** 10.1101/2024.02.16.580709

**Authors:** Michael E. Runda, Hui Miao, Niels A. W. de Kok, Sandy Schmidt

## Abstract

Rieske non-heme iron oxygenases (ROs) are redox enzymes that are essential for microbial biodegradation and natural product synthesis. These enzymes utilize molecular oxygen for oxygenation reactions, making them very useful in applied enzymology due to their broad reaction scope and high selectivities. The mechanism of oxygen activation in ROs involves electron transfers between redox centers of associated protein components, forming an electron transfer chain (ETC). Although the ETC is essential for electron replenishment, it carries the risk of reactive oxygen species (ROS) formation due to electron loss during oxygen activation. Our previous study linked ROS formation to O_2_ uncoupling in the flavin-dependent reductase of the three-component cumene dioxygenase (CDO). In the present study, we extend this finding by investigating the effects of ROS formation on the multi-component CDO system in a cell-free environment. In particular, we focus on the effects of hydrogen peroxide (H_2_O_2_) formation in the presence of a NADH cofactor regeneration system on the efficiency of CDO catalytic efficacy *in vitro.* Based on this, we propose the implementation of hybrid systems with alternative (non-native) redox partners for CDO, which are highly advantageous in terms of reduced H_2_O_2_ formation and increased product formation. The hybrid system consisting of the RO-reductase from phthalate dioxygenase (PDR) and CDO proved to be the most promising for the oxyfunctionalization of indene, showing a 4-fold increase in product formation (20 mM) over 24 h at a 3-fold increase in production rate compared to CDO-WT.

## INTRODUCTION

The oxyfunctionalization of C-H bonds in organic molecules is an essential reaction catalyzed by various enzymes, including peroxygenases or oxygenases.^1–3^ Unlike peroxygenases, which, as their name implies, utilize H_2_O_2_,^4^ the latter class of enzymes is capable of (also) utilizing molecular oxygen to activate unreactive or inert molecular structures for late-stage modifications.^5^ In addition to their outstanding catalytic performance in terms of selectivity and reaction rates, oxygenases have become an indispensable tool in the design of biotechnological applications such as bioremediation,^6^ lignocellulose combustion,^7^ or (biocatalytic) synthesis.^8^

In the search for potential enzyme candidates to perform oxygenation reactions, the class of Rieske non-heme iron oxygenases (ROs) has emerged as a growing family of highly exciting biocatalysts.^9^ Their potential was initially associated mainly with the oxidation of aromatic compounds, which is the first step in microbial biodegradation pathways.^10^ Nowadays, ROs from procaryotic as well as eukaryotic organisms are reported and found to catalyze crucial steps in catabolic and anabolic biological processes.^11, 12^ These findings have driven the search for novel enzymes and contributed to an extensive increase in the reactivities nowadays accessible by characterized ROs. In addition to hydroxylation reactions, the use of ROs in (chemo)enzymatic synthetic routes with the need for desaturation, sulfidation, demethylation of heteroatoms, or C-C bond formation reactions is becoming increasingly conceivable.^13^ However, some common challenges for biocatalyst development still have to be overcome before ROs can be applied as robust and versatile biocatalysts; in particular, challenges related to biocatalyst stability and turnover number have to be addressed.^9^

Like the extensively studied cytochrome P450s protein family, the activity of ROs is based on an electron transfer chain (ETC) that mediates electrons from NAD(P)H to the catalytically active iron (Fe) in the active site of a terminal oxygenase (RO-Oxy).^14^ While the ETC can consist of two or three redox partners, the gateway and the final destination of electrons within ROs is determined by the flavin prosthetic group of a reductase (RO-Red) and the non-heme catalytic iron embedded in the catalytic α-subunit of the RO-Oxy, respectively.^15^ In addition to limiting the boundaries and direction of the electron flow, both redox centers represent a potential site for electron leakage due to unproductive oxygen activation, commonly defined as O_2_ uncoupling (Scheme 1). In this process, electrons are spontaneously and uncoordinatedly transferred to O_2_, forming reactive oxygen species (ROS),^16^ which increase the oxidative pressure on the reaction system and can negatively affect enzyme activities through inhibition or irreversible inactivation.^17^

**Scheme 1.**
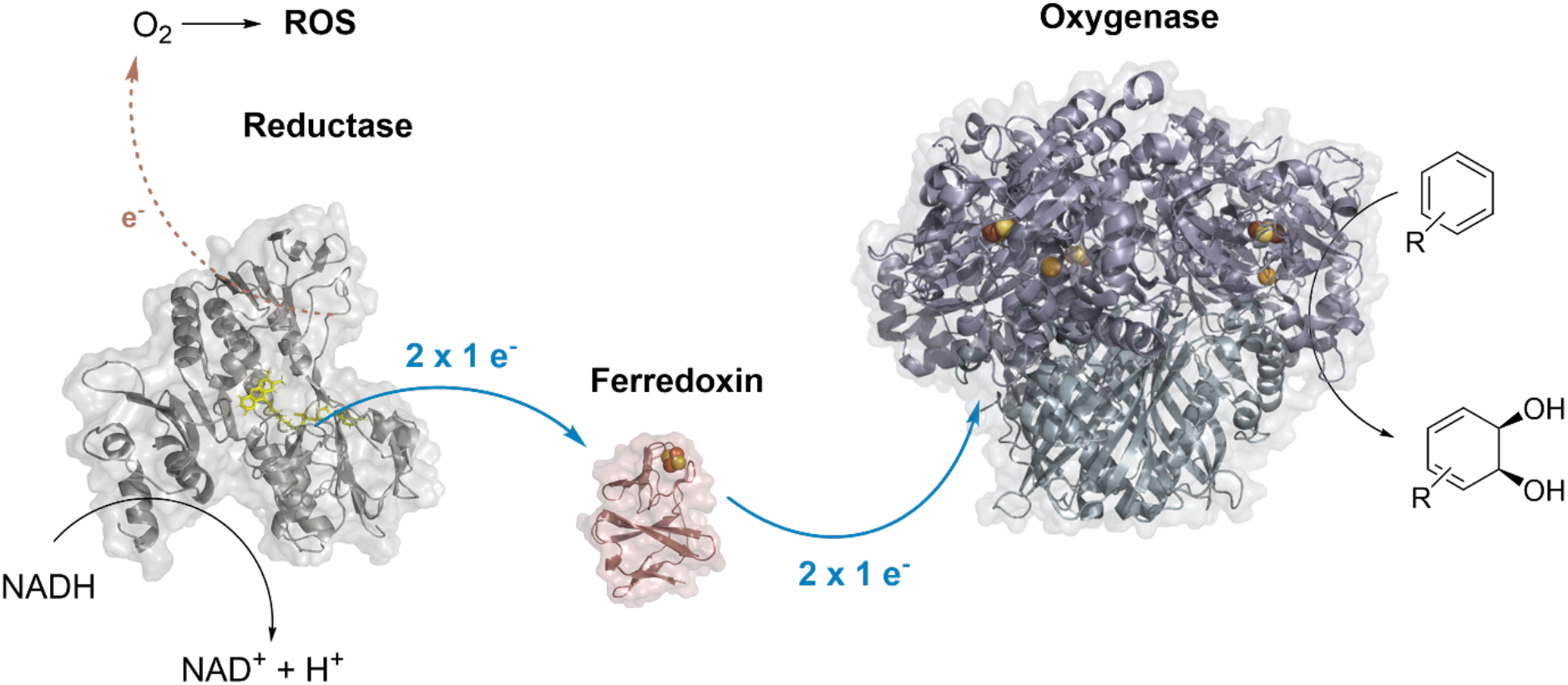
Electron transfer pathway in a typical three-component RO. The dashed arrow highlights O_2_ uncoupling, leading to ROS formation at the reductase component of the RO system. Electron transfer between redox partners required for catalytic activity is highlighted in blue.

Oxygen uncoupling and ROS release in ROs are mainly associated with substrate dependency facilitated by mechanisms occurring at the active site of the RO-Oxy components. As first reported for naphthalene dioxygenase (NDO) from *Pseudomonas sp.* strain NCIB 9816-4, the use of an alternative substrate induces O_2_ uncoupling and the formation of H_2_O_2_ (Equation **1**).^17^

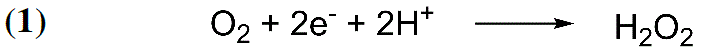

In contrast to the native reactions with naphthalene as substrate, 40-50 % of O_2_ consumed by NDO could be determined as H_2_O_2_ in reactions with benzene. This suggests that H_2_O_2_ acts as an inhibitor and irreversible inactivator of the RO-Oxy of NDO. The same study proposes the occurrence of a Fenton reaction between ferrous Fe and H_2_O_2_ (Equation **2**), resulting in the formation of a highly reactive hydroxyl radical (•OH).^17^ This strong oxidant can react with components of the enzymatic reaction, thus potentially affecting the overall activity of NDO.

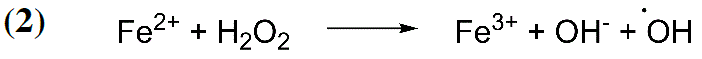

A similar conclusion was drawn from the hydroxylations of various polychlorinated biphenyls (PCBs) catalyzed by biphenyl dioxygenase (BPDO) from *Comamonas testosteroni* B-356.^18^ The coordination of chloro substituents on the substrate influences the effectiveness of O_2_ activation for hydroxylation. While no detectable amounts of H_2_O_2_ were formed in reactions with biphenyl as the substrate, approximately 50 % of O_2_ consumed by PBDO for the conversion of dichlorobiphenyls was reduced to H_2_O_2_. It was proposed that altered substrate coordination within the active site between biphenyl and its substituted analogs resulted in different levels of O_2_ uncoupling.

Recent studies further elucidate the role of substrate-specific O_2_ activation and uncoupling in microbial biodegradation propose substrate-induced O_2_ uncoupling as a potential driving force for ROs to adapt to new contaminants.^16, 19, 20^ Namely, ROS formation due to organic compound exposure could activate an adaptive oxidative stress response and cause an increase in mutation rate.

These findings have contributed significantly to the understanding of the actual reaction mechanism and a possible driving force in the evolution of new ROs.^20^ However, in contrast to the general amount of knowledge on O_2_ uncoupling associated with flavoenzymes,^21^ there is a notable lack of information on this phenomenon in the context of RO-Reds. This is an understudied area of research given the potential impact on the overall catalytic efficiency of the RO system.

One explanation for the lack of reports dealing with the effects of O_2_ uncoupling at RO-Reds may be due to a mechanism that protects O_2_ uncoupling at the flavin prosthetic group. The crystal structure of the three-component toluene dioxygenase (TDO) from *Pseudomonas putida* F1 revealed a stable charge transfer (CT) complex between the reduced RO-Red and NAD^+^. This CT complex presumably suppresses electron transfer from FAD to O_2_.^22^ Conformational changes upon forming the CT complex shield the reactive C4a of the tricyclic isoalloxazine moiety and induce an atypical planar confirmation of FAD, thus, suppressing electron transfer to O_2_. Combined with the rapid electron transfer to the associated ferredoxin, uncoupling at the RO-Red is thought to be prevented under oxic conditions.

The present study evaluates the impact of ROS formation due to O_2_ uncoupling on cumene dioxygenase (CDO) from *Pseudomonas fluorescens* IP01 as a suitable model enzyme system representing a well-studied three-component RO.^23–25^ Therefore, we chose an *in vitro* setup for CDO based on a balanced ratio of RO redox components and a NADH cofactor regeneration system using glucose dehydrogenase (GDH) to maintain cofactor supply.^25^ According to our previous results, the formation of ROS driven solely by RO-Red of CDO (CDO-Red) was demonstrated,^25^ reaching a concentration of more than 4 mM H_2_O_2_ after 24 h in the presence of the NADH-cofactor regeneration system. In addition, the use of catalase was found to be essential to maintain up to 40 % conversion of 10 mM indene (**1**) to 1H-indenol (**1a**) and *cis*-indane diol (**1b**) (Scheme 2). In contrast, reactions without catalase yield in negligible product conversions (< 1 %). These results strongly suggest that the formation of H_2_O_2_ by CDO-Red significantly affects the reaction system, resulting in beneficial effects when a H_2_O_2_ scavenger such as catalase is added.

In addition to removing H_2_O_2_ from the reaction mixture by adding excessive amounts of catalase, a more preventative approach to mitigating ROS formation was sought. As an alternative to enzyme engineering techniques, we used a strategy to replace CDO-Red with other RO-Reds that are less prone to O_2_ uncoupling. The newly created RO systems are composed of redox partners derived from different ROs, herein referred to as hybrid RO systems.

Unexpectedly, we found that RO-Reds from two-component RO systems, such as phthalate dioxygenase (PDR)^26^ or vanillate-*O*-demethylase (VanB),^27^ were able to deliver electrons to the ETC of CDO. This is remarkable because these systems were thought to be incompatible due to the presence of an additional Fe-S cluster domain in these RO-Reds, which replaces the Fd component in those systems. As such, we were able to detect a productive interaction between the RO-Reds and the Fd of CDO (Scheme 2), and an ABTS-HRP assay revealed significantly lower levels of H_2_O_2_ formation when VanB or PDR were used compared to CDO-Red. Therefore, VanB and PDR were promising candidates to form hybrid-CDO *in vitro* systems.

Overall, a significant increase in reaction rate (3-fold) and an excellent maximum product formation of 20 mM was achieved with the hybrid CDO-PDR system compared to the native CDO system. This highlights redox partner exchange as a promising strategy to cope with O_2_ uncoupling in multicomponent enzyme reaction systems.

**Scheme 2.**
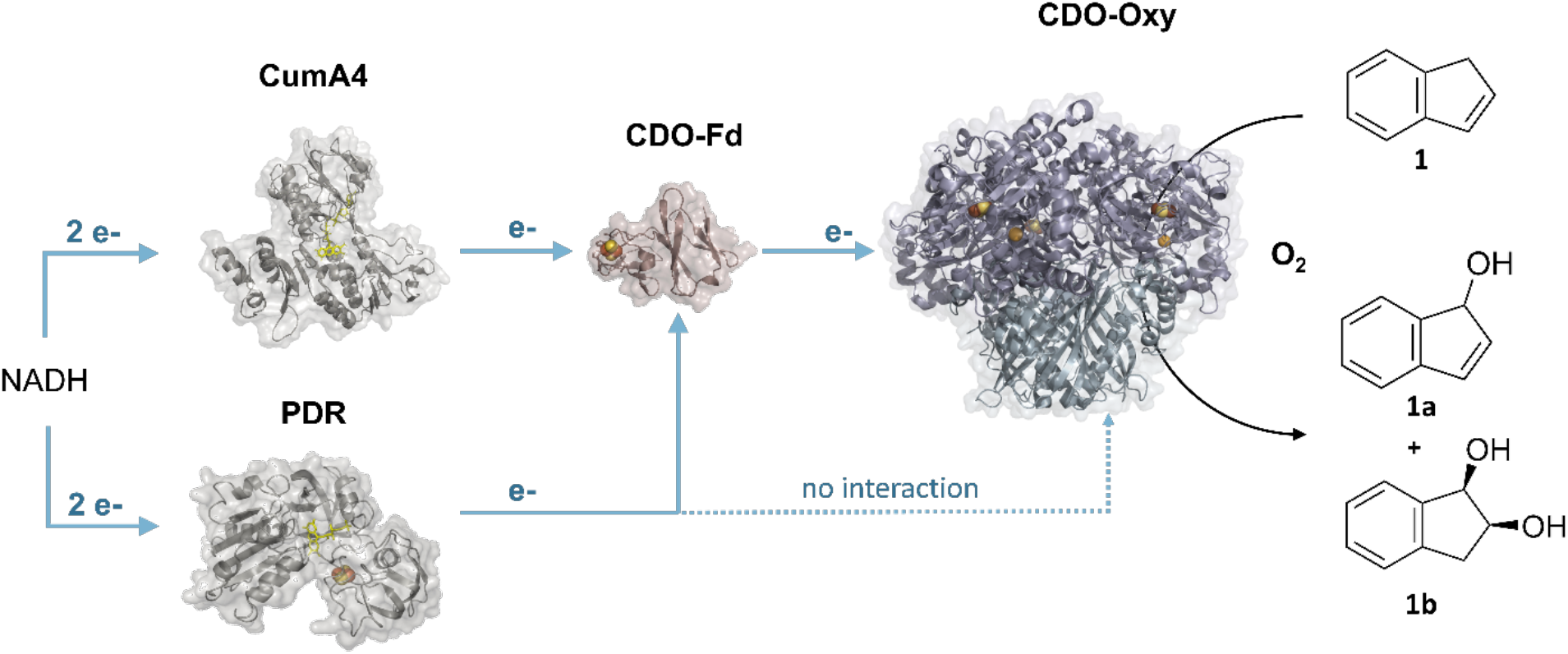
Redox interactions between associated CDO components with PDR (or VanB, not shown) within a hybrid RO system, exemplified for the conversion of indene (**1**) into 1*H*-indenol (**1a**) and *cis*-1,2-indanediol (**1b**) catalyzed by CDO in the presence of molecular oxygen and NADH.^25^

## MATERIALS AND METHODS

### Preparation of *E. coli* JM109(DE3) whole-cell biocatalysts

Resting *E. coli* JM109(DE3) cells containing heterologously expressed RO protein components were applied for whole-cell reactions. Therefore, pUC-based plasmid constructs harboring genes for RO components under the control of a single IPTG (isopropyl ß-D-1-thiogalactopyranoside)– inducible promoter were used (Table S4). Initially, high-cell density precultures were prepared from cryo-glycerol stocks in baffled Erlenmeyer flasks containing LB (1/5th of total flask volume) supplemented with 100 µg mL^-1^ ampicillin. The inoculated pre-cultures were incubated for approximately 16 h (overnight) at 37 °C and 200 rpm prior to inoculation of the main-expression cultures. To ensure proper aeration during cultivation, the main expression cultures were performed in baffled Erlenmeyer flasks filled with TB-media (1/5th of the total flask volume) supplemented with 100 µg mL^-1^ ampicillin. The main cultures were inoculated by adding respective precultures to initial cell densities of OD_600_ 0.1. After incubation at 37 °C and 120 rpm to a cell density between OD_600_ of 0.6 and 0.8, expression was induced by adding 200 µM of IPTG. After induction, the cultures were further incubated for 20 h at 20 °C and 120 rpm. The cells were then harvested by centrifugation for 20 min at 4347x *g* and 5 °C. After washing with 50 mM sodium phosphate buffer (SPB) at a pH of 7.2, a cell suspension with a final concentration of 0.4 g mL^-1^_(*cww*)_ was prepared. The resting cells were kept on ice before usage.

### Protein Expression and Purification for *in vitro* biotransformation studies

Heterologous expression of RO protein components was performed using *E. coli* JM109(DE3) as the expression strain. Aiming for high-level protein expression and the purpose of downstream purification, respective genes were codon-optimized and cloned into pET-28a(+)-based vector backbones in-frame with an N-terminal 6xHis-tag.

Initially, high-cell density precultures were prepared from cryo-glycerol stocks in baffled Erlenmeyer flasks containing LB (1/5th of total flask volume) supplemented with 50 µg mL^-1^ kanamycin. The inoculated pre-cultures were incubated for approximately 16 h (overnight) at 37 °C and 200 rpm prior to inoculation of the main-expression cultures. The main expression cultures were performed in baffled Erlenmeyer flasks filled with TB-media (1/5th of the total flask volume) supplemented with 50 µg mL^-1^ kanamycin, and inoculated by adding the precultures to starting OD_600_ of 0.1. After incubation at 37 °C and 120 rpm to a cell density OD_600_ of about 1.0, expression was induced by adding 50 µM of IPTG. After induction, the cultures were further incubated for 20 h to 22 h at 20 °C and 120 rpm.

Cell harvesting, washing and lysis were performed according to a previously optimized purification protocol.^25^ After incubation of the clarified lysate with 2 mL Ni-sepharose resin (1 column volume (CV) corresponds to 2 mL) for 1 h and washing with 5 CV of washing buffer (50 mM SPB, 30 mM imidazole, 300 mM NaCl, 10 % glycerol, pH 7.2), elution was performed using defined amounts of elution buffer (50 mM SPB, 400 mM imidazole, 300 mM NaCl, 10 % glycerol, pH 7.2). For CDO protein components, elution fractions 0.5 to 2xCVs were collected for downstream sample preparation. For VanB and PDR, elution fractions 1 to 3xCV were collected. To transfer the proteins into the final storage buffer (50 mM SPB, 300 mM NaCl, 10% glycerol, pH 7.2), buffer exchange was performed using PD-10 desalting columns purchased from Cytiva (Danaher, Massachusetts, US). Final protein concentrations were determined using a colorimetric Coomassie protein assay as described previously.^25^

### Sample preparation for *in vitro* reactions

*In vitro* reactions were prepared in 20 mL tightly-sealed glass vials on a 1 mL scale. Unless otherwise stated, a standard reaction solution contained purified protein components at the indicated concentrations, 50 mM glucose, 10 U mL^-1^ glucose dehydrogenase (GDH) from *Bacillus megaterium*, 1 mM dithiothreitol (DTT), 50 mg mL^-1^ catalase from bovine liver, and 400 µM NAD^+^. **1** as substrate was added directly to the reaction solutions prepared in 50 mM SPB (pH 7.2). Reactions were performed in an incubation shaker at 30 °C at 120 rpm for the indicated time. For *in vitro* biotransformations containing 30 mM substrate, DMSO was added as cosolvent to a final concentration of 5 %.

### Sample preparation for whole-cell reactions

Reactions using whole-cell biocatalysts were prepared in 20 mL tightly-sealed glass vials on a 1 mL scale. Therefore, resting *E. coli* JM109(DE3) cells were used as biocatalysts at a final concentration of 0.2 g mL^-1^_(*cww*)_. Standard reaction solutions also contained 50 mM glucose, substrate **1** and 50 mM SPB (pH 7.2) as reaction buffer.

### Quantification of products using GC-FID

Products formed in *in vitro* or whole-cell reactions were quantified via GC-FID using a non-chiral OPTIMA™ 5MS column (Macherey-Nagel GmbH & Co. KG, Düren, Germany). Substrates and products were directly extracted from the aqueous reaction solutions with DCM containing 2 mM acetophenone as an internal standard. Reaction solutions were saturated with NaCl before liquid-liquid extraction to improve phase separation. The organic phase was dried over anhydrous MgSO_4_ before injecting into the GC-FID. The quantitative parameter applied for GC-FID can be found in the Supporting Information (Table S5).

### Quantification of H_2_O_2_ using colorimetric ABTS-HRP Assay

The quantification of H_2_O_2_ in aqueous reaction solutions was performed via a colorimetric ABTS-HRP assay (see reference).^25^

## RESULTS AND DISCUSSION

Driven by our previous findings indicating significant O_2_ uncoupling at the CDO-Red,^25^ we became interested in further evaluating the formation of ROS in the native reaction system. While the term ROS encompasses a class of oxygen-derived moieties resulting from single-electron reduction steps, we have a strong indication that the impact of H_2_O_2_ on the CDO system is particularly critical. This assumption is based on the previously reported necessity of catalase to establish a suitable *in vitro* system for CDO.^25^ In addition, the availability of easy-to-use quantitative assays at the microtiter plate scale further convinced us to consider H_2_O_2_ as a representative and valid ROS in our study. Initially, we were interested in H_2_O_2_ formation at CDO-Red in terms of cofactor availability. The activities of ROs are dependent on electrons from the reduced nicotinamide cofactors NAD(P)H. Efforts to circumvent the need for stoichiometric amounts of NAD(P)H have led to promising and effective cofactor regeneration systems.^28^ While the use of a GDH-based cofactor regeneration system is successful in maintaining a sufficient amount of NADH for the functionalization of **1** with CDO in an optimized reaction setup, the question remains as to how it affects H_2_O_2_ formation. To address this question, *in vitro* reactions were performed using CDO-Red together with the GDH-based cofactor regeneration system (Figure 1a, Scheme 3). The amount of H_2_O_2_ detected after 24 h exceeded 4 mM in the reaction mixture. Compared to our previously reported results, where 35 µM H_2_O_2_ was formed in the reaction with reduced NADH cofactor,^25^ this value represents a more than 100-fold increase of H_2_O_2_ and thus contributes substantially to the oxidative pressure of the reaction environment.

**Figure 1.**
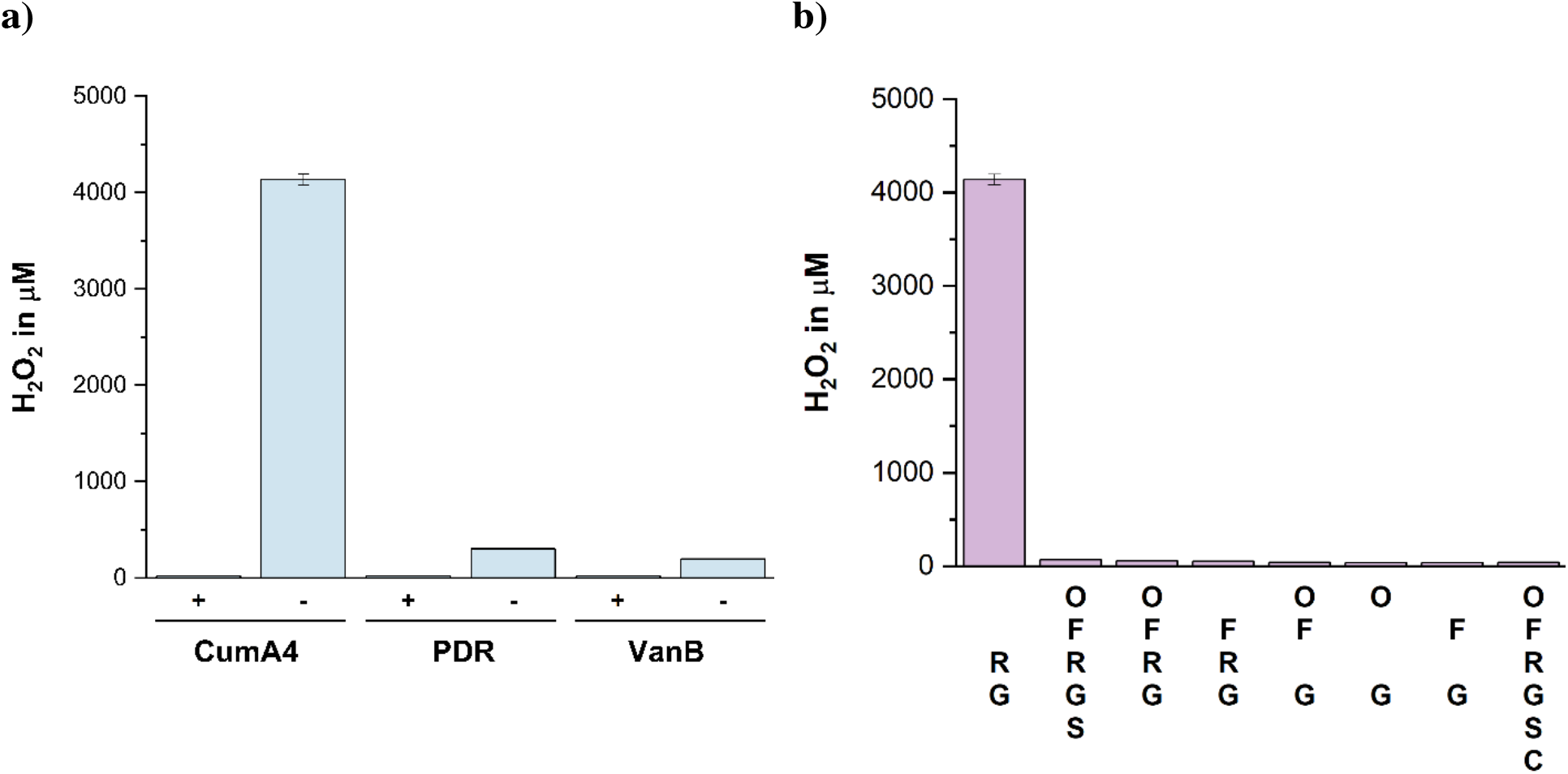
ROS formation driven by O_2_ uncoupling at RO-Reds in *in vitro* reactions. **a)** Total concentrations of H_2_O_2_ determined after 24 h in reactions with indicated RO-Reds in the presence (+) or absence (-) of catalase. **b)** Total concentrations of H_2_O_2_ determined after 24 h in *in vitro* reactions with indicated reaction components: O: CDO-Oxy; F: CDO-Fd; R: CDO-Red; G: GDH-based NADH-regeneration system; S: substrate; C: catalase.

As reported for TDO, RO-Red forms a stable complex with oxidized NAD^+^ during electron transfer, suppressing the reaction between FAD and O_2_ under oxic conditions and thus preventing O_2_ uncoupling.^22^ We infer that by implementing of a cofactor regeneration system, as used in this study, kept the concentration of oxidized NAD^+^ in the reaction solutions low and thus interfered with the formation of the CT complex at CDO-Red. Together with the absence of associated redox centers to accept electrons from the flavin prosthetic group, a significant increase in H_2_O_2_ due to O_2_ uncoupling was expected. In addition, the H_2_O_2_ formation by PDR and VanB was evaluated analogously. The amount of H_2_O_2_ formed by PDR and VanB over 24 h was significantly lower than that formed by CDO-Red, with values of approximately 300 and 200 µM H_2_O_2_ for PDR and VanB, respectively (Figure 1a).

In contrast to CDO-Red, PDR and VanB are derived from two-component ROs and have a prominent Fd domain containing a [2Fe-2S] cluster.^26, 29^ It is proposed that electrons stored in the fully reduced state of the flavin prosthetic group are successively transferred intramolecularly to the [2Fe-2S] redox center. From there, these electrons are transferred to an oxidized Fe-S cluster within a Rieske center of an associated RO-Oxy. We hypothesize that the additional Fe-S cluster in the protein scaffold of RO-Reds reduces the electron loss at the flavin prosthetic group and thus has a beneficial effect on the present *in vitro* system with respect to H_2_O_2_ formation (Scheme 3).

**Scheme 3.**
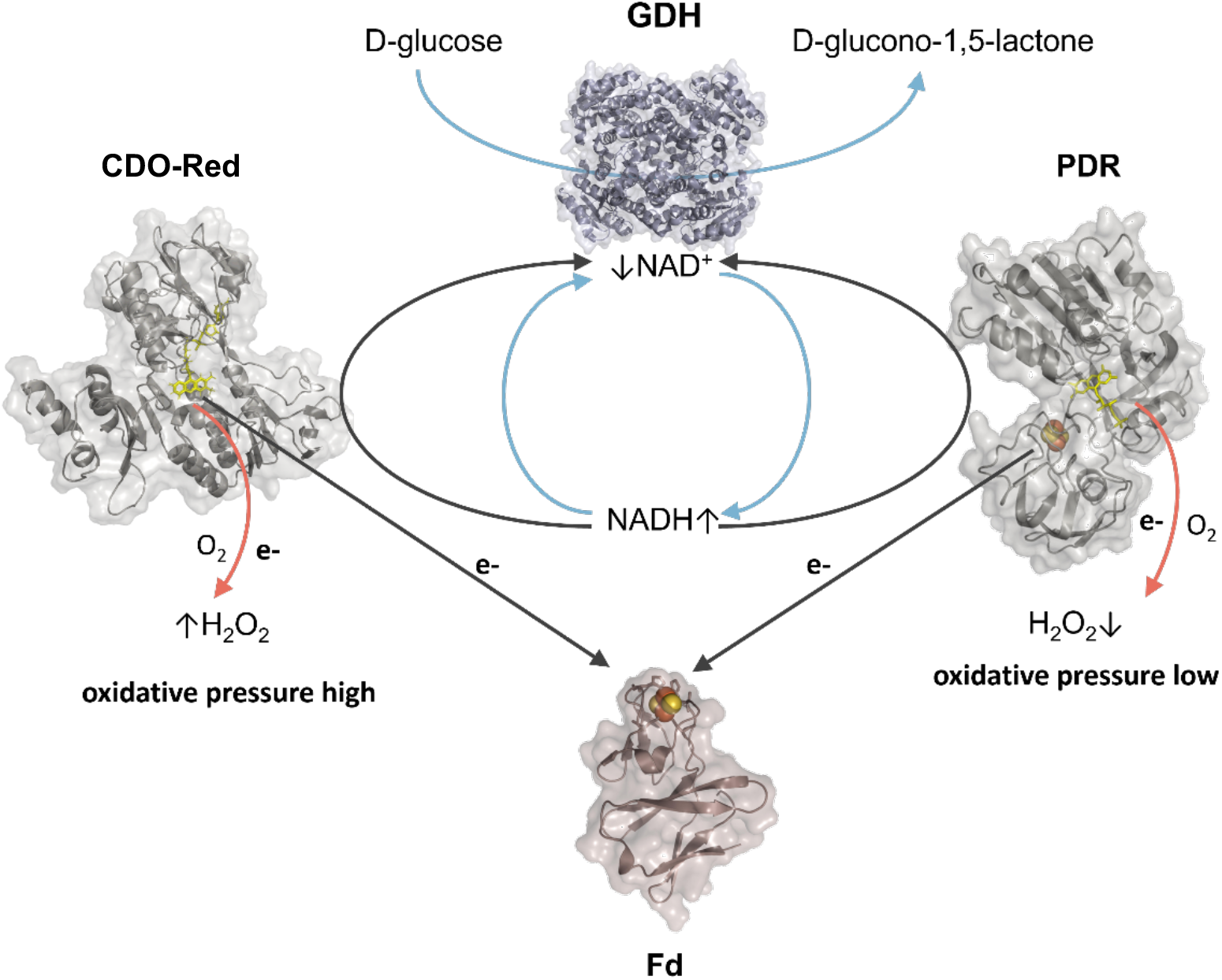
Schematic representation of O_2_ uncoupling and H_2_O_2_ formation in CDO-Red and PDR, driven by a high concentration of reduced cofactor due to the NADH regeneration system. Reactions involved in the cofactor regeneration system are highlighted in blue. Flavin prosthetic groups in RO-Reds are shown as yellow sticks in the corresponding 3D models. The [2Fe-2S]-clusters in PDR and CDO-Fd are shown as orange-yellow spheres. Orange arrows indicate the site of potential O_2_ uncoupling leading to the formation of H_2_O_2_.

This observation agrees well with results obtained in control experiments using CDO-Red in the presence of CDO-Fd, its native redox partner in the CDO reaction system (Figure 1b). This indicates that the presence of CDO-Fd has a protective effect on O_2_ uncoupling. As mentioned above, VanB and PDR contain a Fe-S cluster domain; presumably, the presence of this domain protects against O_2_ uncoupling. Furthermore, CDO-Fd and CDO-Oxy do not significantly affect the total ROS formation as confirmed in the ABTS-HRP assay. Therefore, *in situ* formation of H_2_O_2_ driven solely by the NADH regeneration system can be excluded.

Since these results indicate a difference in ROS formation depending on the corresponding RO-Red, we sought to take advantage of this observation. Besides considering taking advantage of CDO-Red’s stability in relatively high H_2_O_2_ concentrations as a potential system for *in situ* H_2_O_2_ generation, we were particularly interested in establishing hybrid systems between the described RO redox components. In addition to gaining valuable insight into the interaction compatibilities between redox partners of two– and three-component RO reaction systems, low oxidative pressure due to Fe-S-containing RO-Reds is considered advantageous for achieving higher catalytic efficiencies. Therefore, we performed *in vitro* reactions for 24 h at 30 °C using optimized conditions previously reported for CDO to investigate component compatibility between different RO systems and to form active hybrid systems.^25^ Interestingly, product quantification via GC-FID analysis indicates the functional redox interaction between VanB and PDR with the Fd component of the CDO, allowing electron transfer from the respective Red to the terminal CDO-Oxy (Figure 2).

**Figure 2.**
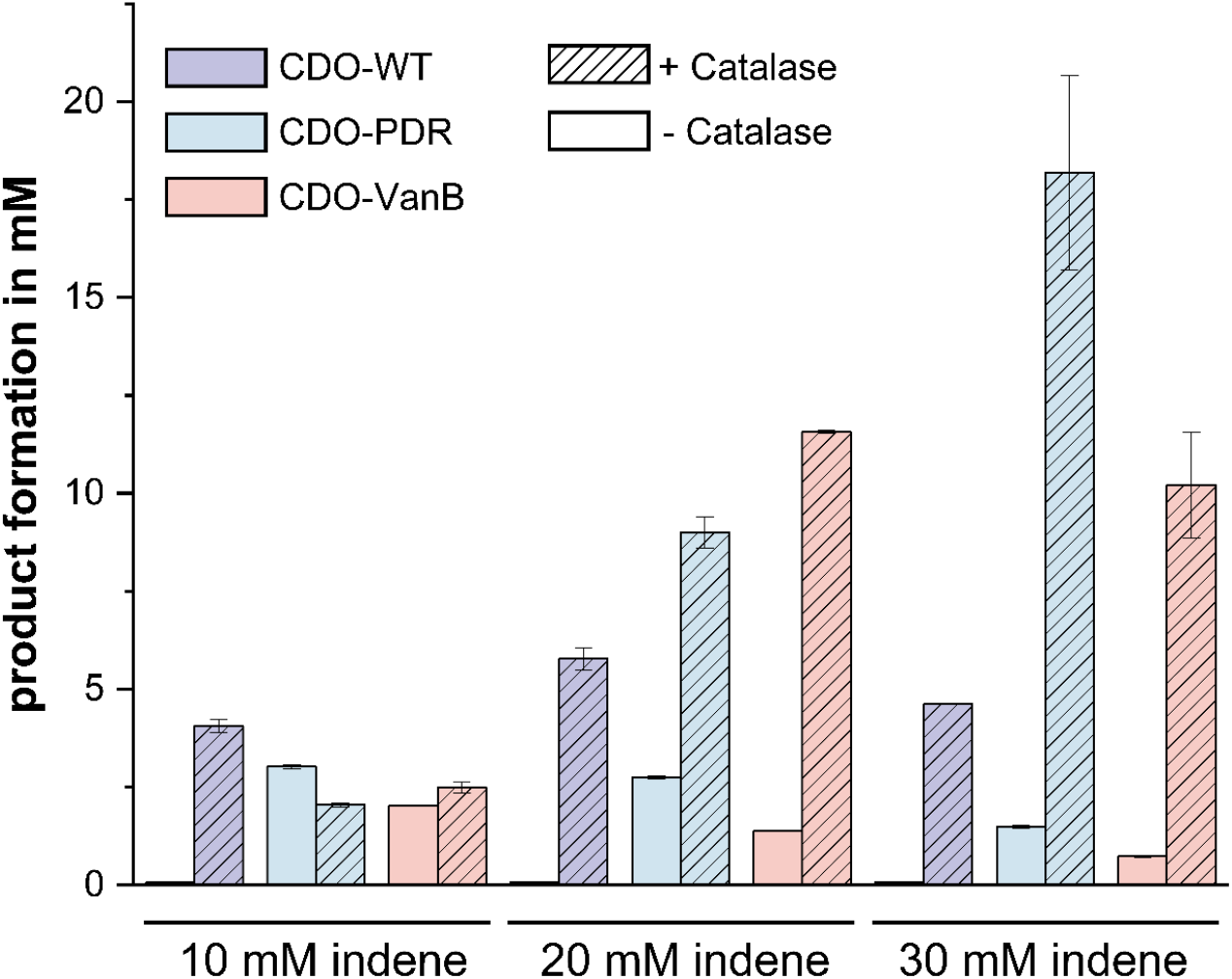
Comparison of product concentrations determined after 24 h in *in vitro* reactions using alternative CDO-PDR and CDO-VanB hybrid systems with CDO-WT. Reactions were performed at different concentrations of **1** and in the presence (+) or absence (-) of catalase. Product formation corresponds to the total amount of **1a** and **1b**.

This observation was rather unexpected since the associated redox partners for VanB and PDR within their native ETC are RO-Oxys, and an RO-Fd is not present in the electron transfer chain of these two-component systems. However, no product formation could be detected in hybrid reactions without the Fd component of CDO (data not shown), leading to the conclusion that less interaction specificity is required between a two-component RO-Red and a three-component RO-Fd.

In addition, H_2_O_2_ formation due to O_2_ uncoupling and the resulting oxidative pressure appears to have a more severe impact on the *in vitro* reaction system for ROs than initially expected. Previous results from the ABTS-HRP assay indicated no insignificant amounts of residual H_2_O_2_ produced by CDO-Red in the presence of CDO-Fd after 24 h (Figure 1b). Moreover, product formation in *in vitro* reactions with complete CDO-WT appears to be strongly dependent on the presence of catalase. Conversely, catalase has less effect on product formation obtained from the PDR– and VanB-CDO hybrid systems, which were previously thought to be less susceptible to O_2_ uncoupling driving ROS formation. Not only does the use of alternative RO-Reds outcompete the CDO-WT system in the absence of catalase as H_2_O_2_ scavenger in the *in vitro* oxyfunctionalization of **1**. The use of alternative RO-Reds also enables product formation of up to 20 mM in the PDR-CDO hybrid system which corresponds to a 4-fold improvement in conversion compared to CDO-WT. In particular, concerning substrate conversion, the product formation catalyzed by CDO-WT appears to be independent of substrate concentrations exceeding 10 mM in the presence of catalase. On the other hand, product formation observed in the reactions using the PDR-CDO hybrid system can be enhanced by increasing the concentration of **1** up to 30 mM. In addition to ruling out inhibitory effects on the CDO-Oxy or substrate limitation, the observed results suggest that one or more reaction components are affected by increased oxidative pressure over time, which could cause the ETC to collapse. Thus, the addition of fresh RO components to the CDO-WT system after 2 h could increase the total product formation up to 3-fold (Figure 3).

**Figure 3.**
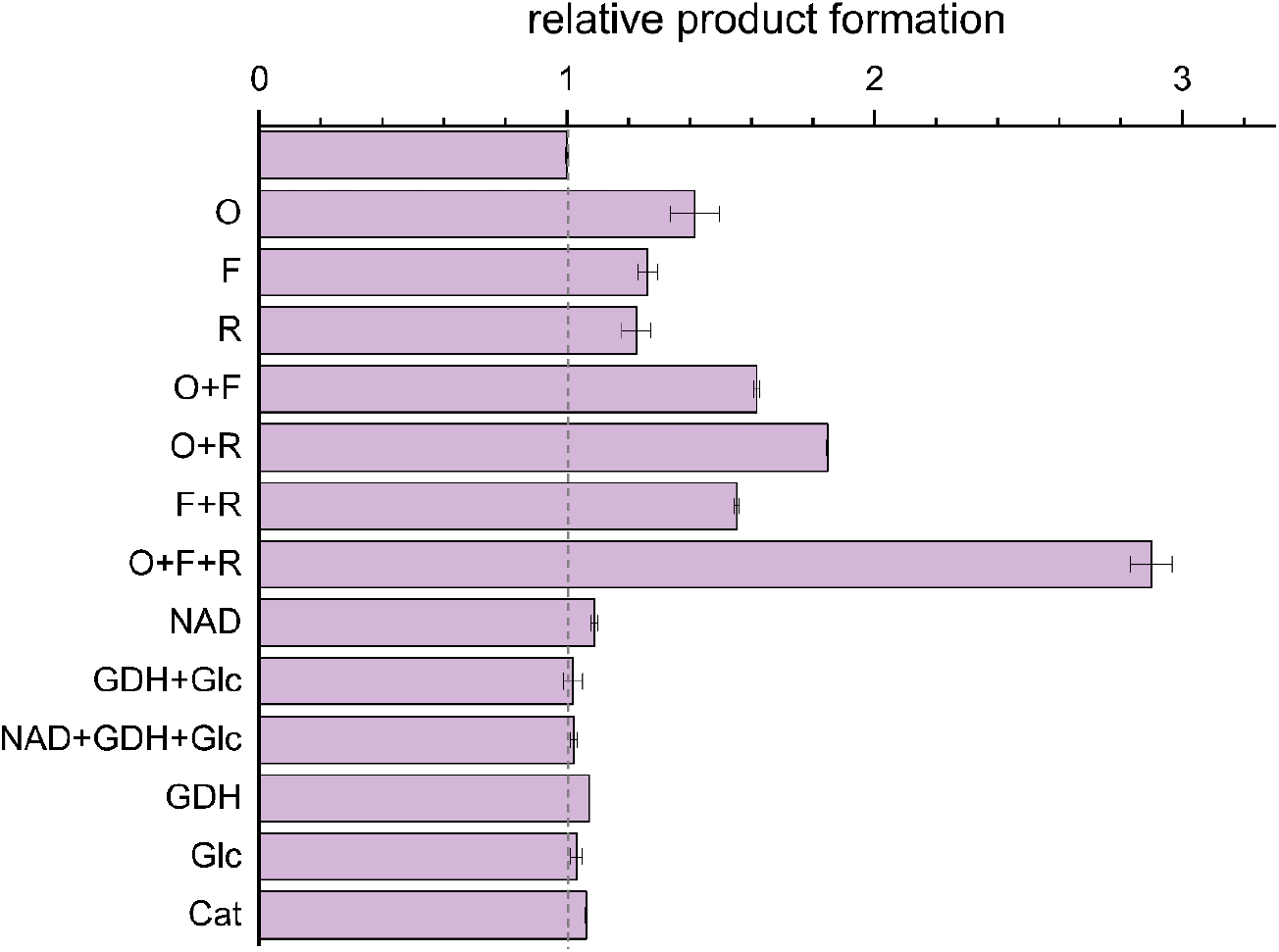
Relative product formation determined in *in vitro* reactions of 30 mM **1** with CDO-WT after 24 h and addition of the indicated reaction components after 120 min: O: CDO-Oxy; F: CDO-Fd, R: CDO-Red; GDH: glucose dehydrogenase; Glc: glucose; Cat: catalase. The value of 1 corresponds to the product formation determined in reactions without the addition of reaction components after 120 min (first bar). Product formation corresponds to the total amount of **1a** and **1b**. The relative product formation of 1 is highlighted as a dashed line.

While the addition of individual protein components has a small effect on the relative conversion of **1**, the improvement in product formation due to synergistic effects outweighs this effect. Based on these results, an exhaustion of the applied cofactor regeneration system or an inactivation of catalase can be excluded. These results lead to the need to consider the ETC as a single entity and its functionality as a product of the activities and conditions of each redox component at a given time. Taking the general concerns regarding the enzyme stability of ROs in cell-free environments into account, the depletion of the ETC might be progressive rather than a defined end-point event. This hypothesis is supported by an overall 3-fold increase in initial rates using the hybrid CDO-PDR *in vitro* system compared to the CDO-WT with values of 95 mU and 35 mU, respectively (Figure S6). Convinced that hybrid systems could be used to establish more efficient and robust *in vitro* reaction platforms for CDO, we sought to evaluate their potential for whole-cell applications. Traditionally, the use of resting *E. coli* cells as biocatalysts for RO-catalyzed oxygenation reactions has proven to be the most suitable in terms of industrial applicability. However, the advantage of using CDO hybrid systems observed with purified enzyme components could not be realized in whole-cell reactions (Figure 4). On the contrary, the exchange of RO-Red seems to be detrimental to the overall product formation.

**Figure 4.**
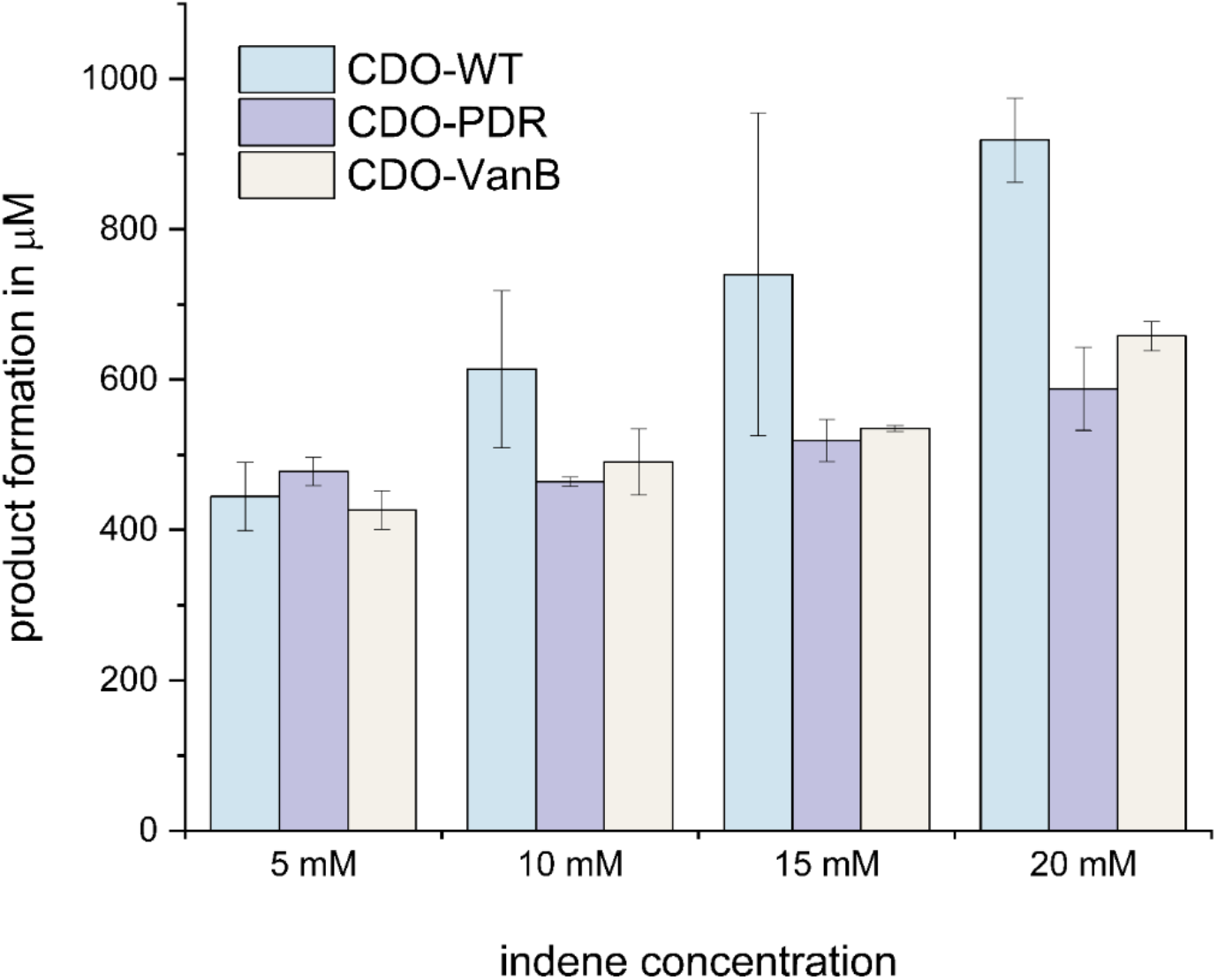
Comparison of product concentrations determined after 24 h in whole-cell reactions obtained with CDO-WT and the proposed CDO-PDR or CDO-VanB hybrid RO-systems. Reactions were performed at different substrate concentrations as indicated. Product formation corresponds to the total amount of **1a** and **1b**.

We assume that this behavior is due to suboptimal expression levels in the host organism and that prevention of ROS formation is less critical. The overexpression of another Fe-S-containing protein, in addition to CDO-Fd and CDO-Oxy, places an additional metabolic burden on *E. coli* and thus negatively affects the overall expression levels of the ROS component. Moreover, considering the protective mechanisms of the cells as part of the intracellular oxidative stress response; the effect of ROS formation, on which the idea of using alternative redox partners is based, is probably less dominant.

In conclusion, this study elaborates on the formation of H_2_O_2_ due to O_2_ uncoupling at RO-Reds and its impact on an *in vitro* reaction setup for the three-component CDO. To design an economically feasible and reliable reaction setup for CDO by implementing a cofactor regeneration system, it was found that the regeneration system significantly contributed to O_2_ uncoupling. As such, the continuous cofactor regeneration interferes with the formation of stable CT complexes between reduced CDO-Red and NAD^+^, which are reported to be crucial for TDO. To cope with O_2_ uncoupling in the presence of a cofactor regeneration system, the possibility of using alternative RO Reds, which are less prone to *in situ* H_2_O_2_ generation, was investigated.

Reductases containing [2Fe-2S], such as VanB and PDR, were found to significantly reduce O_2_ uncoupling and thus overall H_2_O_2_ formation in the *in vitro* reaction system. Furthermore, we established hybrid CDO systems using non-native Reds such as VanB and PDR derived from two-component RO systems and demonstrated that their interaction with CDO-Fd is productive to efficiently promote the conversion of **1**. Overall, the PDR-CDO hybrid system was most promising in terms of maximum product formation and activity, outperforming the CDO-WT reaction system.

In summary, this report describes a strategy to optimize multi-component enzyme redox systems suffering from O_2_ uncoupling by exploiting redox partner compatibility. Furthermore, the reported redox partner flexibility between two– and three-component ROs provides exciting insights for further studies to elucidate redox interactions and the specificity required for activity.

## ACCESSION CODES

- CDO-Oxy α-subunit CumA1: Q51743
- CDO-Oxy β-subunit CumA2: Q51744
- CDO-Fd CumA3: Q51746
- CDO-Red CumA4: Q51747
- VanB: Q9HUQ8
- PDR: P33164

## ASSOCIATED CONTENT

### Supporting Information

The Supporting Information includes additional experimental procedures, including a list of bacterial strains and plasmids used, the cloning, heterologous expression, and purification of CDO and reductases; procedures for *in vitro* and whole-cell biotransformations, GC-FID product analysis, ABTS-HRP assay; Tables S1-S5; and Figures S1−S6.

## AUTHOR INFORMATION

### Corresponding Author

Sandy Schmidt − Department of Chemical and Pharmaceutical Biology, University of Groningen, Groningen Research Institute of Pharmacy, Antonius Deusinglaan 1, 9713AV Groningen, The Netherlands; orcid.org/0000-0002-8443-8805;

### Authors

**Michael E. Runda** − Department of Chemical and Pharmaceutical Biology, University of Groningen, Groningen Research Institute of Pharmacy, Antonius Deusinglaan 1, 9713AV Groningen, The Netherlands; orcid.org/0000-0002-2569-8260

**Hui Miao** − Department of Chemical and Pharmaceutical Biology, University of Groningen, Groningen Research Institute of Pharmacy, Antonius Deusinglaan 1, 9713AV Groningen, The Netherlands; orcid.org/0009-0003-6765-9766

**Niels A. W. de Kok** − Department of Chemical and Pharmaceutical Biology, University of Groningen, Groningen Research Institute of Pharmacy, Antonius Deusinglaan 1, 9713AV Groningen, The Netherlands; orcid.org/0000-0002-0306-892X

### Author Contributions

All authors (M.E.R., H.M., N.A.W.d.K. and S.S.) contributed to the design of the experiments and wrote the manuscript. M.E.R. performed cloning experiments, purified all proteins used in this work, and performed all activity measurements. H.M. contributed to data acquisition and sample preparation for *in vitro* and whole-cell reactions. The manuscript draft was written by M.E.R. and all authors contributed to writing and reviewing. All authors have approved the final version of the manuscript.

### Notes

The authors declare no competing financial interest.

## ABBREVIATIONS

ABTS, 2,2’-azino-bis(3-ethylbenzothiazoline-6-sulfonic acid); biphenyl dioxygenase, BPDO; CDO, cumene dioxygenase; CT, charge transfer; *cww*, cell wet weight; DTT, dithiothreitol; Fd, ferredoxin; H_2_O_2_, hydrogen peroxide; IPTG, isopropyl ß-D-1-thiogalactopyranoside; LB, lysogeny broth; Oxy, terminal RO oxygenase; polychlorinated biphenyl, PCB; Red, reductase; RO, Rieske oxygenase; ROS, reactive oxygen species; SPB, sodium phosphate buffer; TB, terrific broth; TDO, toluene dioxygenase

