## Supplementary Material for "Developing Hybrid Systems to Address O_2_ Uncoupling in Multi-Component Rieske Oxygenases"

### List of Abbreviations

| | |
|---|---|
| ABTS | 2,2'-azino-bis(3-ethylbenzothiazoline-6-sulfonic acid |
| CDO | cumene dioxygenase |
| c.o. | codon-optimized |
| GC | gas chromatography |
| FID | flame ionization detector |
| GDH | glucose dehydrogenase |
| HRP | horseradish peroxidase |
| IPA | integrated peak area |
| IST | internal standard |
| ORF | open reading frame |
| RO | Rieske oxygenase |
| RT | retention time |
| RRF | relative response factor |

### Materials

Unless indicated otherwise, all chemicals and solvents were purchased from Sigma-Aldrich (St. Louis, USA), Merck KGaA (Darmstadt, Germany), TCI- Tokyo Chemical Industry (Tokyo, Japan), Carl Roth GmbH + Co. KG (Karlsruhe, Germany) and Duchefa Biochemie (Haarlem, The Netherlands) in the highest purity available. Enzymes for Gibson Assembly were purchased from New England Biolabs (Ipswich, Massachusetts, US). FastDigest™ enzymes from Thermo Scientific (Massachusetts, USA) were used for restriction digestion of DNA. Kits for PCR product purification, DNA gel extraction, and plasmid isolation were used from Qiagen or MACHEREY-NAGEL (Düren, Germany). The service of Macrogen Europe (Amsterdam, The Netherlands) was used for DNA sequencing. Synthetic genes were purchased from TWIST Bioscience (South San Francisco, California, US). GC-FID analyses were performed on a Shimadzu GC-2010 Plus equipped with an AOC-20i auto-injector. Column specifications and conditions for GC analyses are detailed in another section of the Supporting Information.

### Bacterial Strains

During the project, different *E. coli* strains were used for cloning, protein expression, and as biocatalysts in whole-cell reactions. Used *E. coli* strains are summarized in **Table S1**.

**Table S1. *E. coli* strains used for molecular cloning, protein expression, and as whole-cell biocatalysts.**

| Strain | Genotype | Description/Use |
|---|---|---|
| **DH5α** | $F^-$ ϕ80*lac*ZΔM15 Δ(*lac*ZYA-*arg*F)U169 *rec*A1 *end*A1 *hsd*R17($rK^-$, $mK^+$) *pho*A *sup*E44 $\lambda^-$*thi*-1 *gyr*A96 *rel*A1 | routine cloning |
| **JM109 (DE3)** | *end*A1, *rec*A1, *gyr*A96, *thi, hsd*R17 ($r_k^-$, $m_k^+$), *rel*A1,*sup*E44, λ–, Δ(*lac-pro*AB), [F′, *tra*D36, *pro*AB, *lac*$I^q$ZΔM15], lDE3 | expression host for RO protein components, whole-cell biocatalysts |
| **BL21(DE3)** | $F^-$*omp*T *hsdS$_B$* ($r_B^-$, $m_B^-$) *gal dcm* (DE3) | expression host for GDH |

### Plasmid Constructs

#### Plasmid constructs for the expression and purification of CDO

Plasmid constructs applied for the heterologous expression of CDO protein components in *E. coli* JM109(DE3) were designed and reported in a previous study (see **Table S4**).[1]

#### Plasmid constructs for the expression and purification of VanB and PDR

Plasmid constructs for the expression of VanB and PDR were created via restriction-ligation cloning. For the purpose of downstream protein purification using affinity chromatography, the relevant genes were cloned in-frame with a 6xHis-tag of pET-28a(+). Therefore, *BamH*I and *Xho*I restriction sites were chosen as most suitable for assembling inserts and linearized vector backbones.

Gene fragments for VanB and PDR with *BamH*I and *Xho*I restriction sites attached, were synthesized by TWIST Bioscience (South San Francisco, California, US). Prior to synthesis, genes were optimized for the heterologous expression in *E. coli* using the supplier's algorithm.

Oligonucleotides used for the DNA amplification during restriction cloning are indicated in **Table S2**.

The success of the Restriction cloning was evaluated by DNA sequencing.

**Table S2. Oligonucleotides used for restriction cloning.**

| Name | Sequence 5'→ 3' | Description |
|---|---|---|
| MRU_130 | GCCGCACTCGAGCACC | Forward primer used for PCR amplification of pET-28a(+) vector backbone. |
| MRU_85 | GCAAGCTTGTCGACGGAG | Reverse primer used for PCR amplification of pET-28a(+) vector backbone. |
| TWIST_F* | CTAGTTATTGCTCAGCGGT | Forward primer used for the amplification of gene fragments synthesized by TWIST Bioscience |
| TWIST_R* | GCCAATCTCCTCGGGACTT TGC | Reverse primer used for the amplification of gene fragments synthesized by TWIST Bioscience |

*Oligonucleotides bind to flanking regions which, by default, are attached to synthesized DNA fragments purchased from TWIST Bioscience.

#### Plasmid constructs for the co-expression of CDO-hybrid systems

For the preparation of resting *E. coli* cells as biocatalysts for testing the proposed CDO-hybrid systems in whole-cells, plasmid constructs were created. Thereby, the gene encoding for CumA4 within the ORF at the parental pIP107D plasmid (see **Table S4**) was exchanged with codon-optimized (c.o.) genes for VanB or PDR, respectively. Due to the lack of suitable restriction sites on pIP107D, Gibson Assembly was performed. The backbone fragment used in Gibson Assembly was obtained by PCR amplification of pIP107D. The associated inserts with the compatible Gibson 5'overhangs attached were obtained via PCR using pET28a-[N-His]-VanB or pET28a-[N-His]-PDR as templates (see **Table S4**). After *Dpn*I digestion and gel extractions, purified DNA fragments were added at an equimolar ratio (total volume 6 µL) to a 15 µL aliquot of Gibson Master Mix. After incubation for 1 h at 50 °C, the reaction mixture was directly used for transformation into chemically competent *E. coli* DH5α cells. The success of the Gibson Assembly was evaluated by colony PCR followed by DNA sequencing.

Oligonucleotides used for the amplifications of pET-28a(+) and Gibson inserts are summarized in **Table S3.**

**Table S3. Oligonucleotides used for Gibson Assembly.**

| Name | Sequence 5'→ 3' | Description |
|---|---|---|
| MRU_186_pCDO_lin_F | GGCGCTAGATACCCGGCATC | Forward primer used for PCR amplification of pET-28a(+) vector backbone. |
| MRU_187_pCDO_lin_R | CGAGCTCGAATTCACTGGCC | Reverse primer used for PCR amplification of pET-28a(+) vector backbone. |
| MRU_190_VanB-Gibson_F | GTGTTCGTAGACTTTGATGCCGGGTATCTAGCGCCATGATCGAAGTTATTGTTGGTGCC | Forward Gibson primer used for PCR amplification of VanB |
| MRU_191_VanB-Gibson_R | ACGTTGTAAAACGACGGCCAGTGAATTCGAGCTCGTTACAGATCCAGAACCAGACGC | Reverse Gibson primer used for PCR amplification of VanB |
| MRU_188_PDR-Gibson_F | GTGTTCGTAGACTTTGATGCCGGGTATCTAGCGCCATGACCACACCGCAAGAAGA* | Forward Gibson primer used for PCR amplification of PDR |
| MRU_189_PDR-Gibson_R | ACGTTGTAAAACGACGGCCAGTGAATTCGAGCTCGTTACAGATCCAGAACCAGTTCTGC* | Reverse Gibson primer used for PCR amplification of PDR |

*Binding region highlighted in grey.

**Table S4. Plasmid constructs used for expression of RO proteins in *E. coli*.**

| Construct | Vector | Insert |
|---|---|---|
| pET28a-[N-His]-CumA1-CumA2 | pET-28a(+) | *cumA1* (N-terminal 6xHis-tag, c.o.), *cumA2* (c.o.) |
| pET28a-[N-His]-CumA3 | pET-28a(+) | *cumA3* (N-terminal 6xHis-tag, c.o.) |
| pET28a-[N-His]-CumA4 | pET-28a(+) | *cumA4* (N-terminal 6xHis-tag, c.o.) |
| pET28a-[N-His]-VanB | pET-28a(+) | *vanB* (N-terminal 6xHis-tag, c.o.) |
| pET28a-[N-His]-PDR | pET-28a(+) | pdr (N-terminal 6xHis-tag, c.o.) |
| pIP107D | pUC | *cumA1, cumA2, cumA3, cumA4* |
| pIP107D-VanB | pUC | *cumA1, cumA2, cumA3, vanB* |
| pIP107D-PDR | pUC | *cumA1, cumA2, cumA3, pdr* |

### Analytics

#### GC Analysis

##### Parameters applied in non-chiral GC-FID analysis

Parameters applied for non-chial GC-FID analysis are summarized in **Table S5.**

**Table S5. GC-FID parameters applied in this study**

| | |
|---|---|
| Gas chromatograph | Shimadzu GC-2010 Plus (Kyoto, Japan) |
| Column | Optima™ 5 MS GC from Macherey-Nagel™ |
| length<br>inner diameter<br>film thickness | 30 m<br>0.25 mm<br>0.25 µM |
| Injection volume | 1 µL |
| Injection temp. | 230 °C |
| Injection mode | Split |
| Carrier gas | $N_2$ |
| Flow control mode | Linear velocity |
| Pressure | 116.3 kPa |
| Total flow | 54.0 mL $min^{-1}$ |
| Column flow | 2.37 mL $min^{-1}$ |
| Linear velocity | 49.1 cm $s^{-1}$ |
| Purge flow | 3.0 mL $min^{-1}$ |
| Split ratio | 20.5 |
| Oven temp. program | 70 °C, 5 °C $min^{-1}$ to 110 °C, 15 °C $min^{-1}$ to 280 °C, hold for 5 min |
| FID temperature | 250 °C |
| Retention times | indene: 5.90 min<br>acetophenone (IS): 6.27 min<br>1*H*-indenol: 9.58 min<br>*cis*-indanediol: 12.50 min |

##### Calibration of GC-FID for product quantification

Determining the concentrations of total product was conducted using linear regression of established calibration curves (**Figures S1** and **S2**). For accurate results, relevant compounds were extracted from aqueous solutions, following the protocol indicated in the section '*Quantification of products using GC-*

*FID'* indicated in the method section of the main manuscript. In addition, acetophenone was used as an internal standard (2 mM).

*Calibration of 1H-indenol*

The product 1*H*-indenol could not be used directly for calibration as this compound is not commercially available and the synthesis was reported as challenging due to the formation of explosives.[2]

Because of that, we first established a calibration curve for 1-indanone (see **Figure S1**), which was then used to evaluate the concentration of 1*H*-indenol by approximation via the relative response factor (RRF).[1]

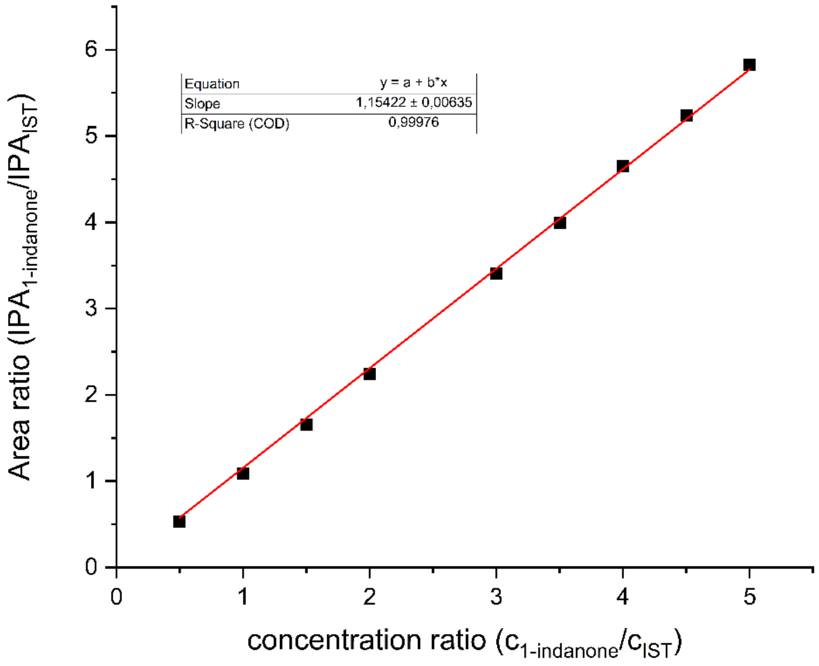

**Figure S1.** Calibration curve for 1-indanone using acetophenone as IST with non-chrial GC-FID. Applied GC parameters can be found in Table S5. The linear regression is indicated.

*Calibration of cis-indanediol*

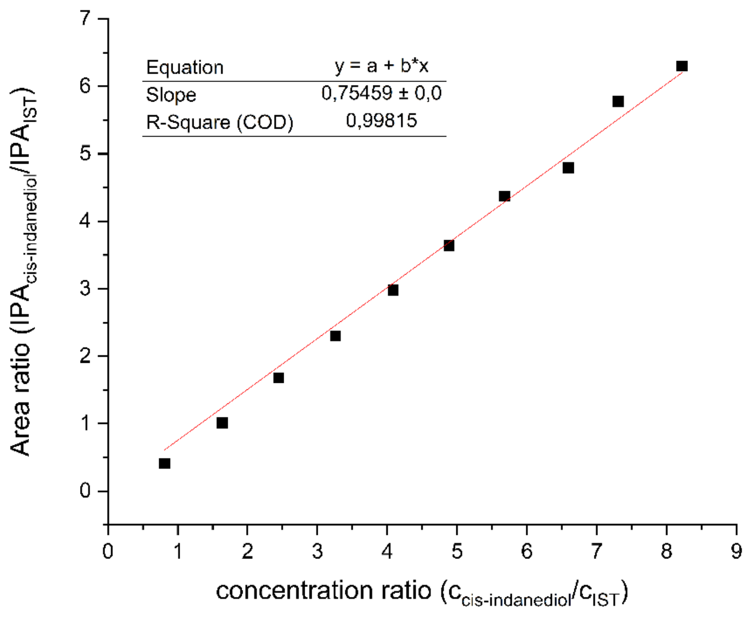

**Figure S2.** Calibration curve for *cis*-indanediol using acetophenone as IST with non-chiral GC-FID. Applied GC parameters can be found in **Table S5**. The linear regression is indicated.

#### Example chromatograms obtained from GC-FID analyses

Representative extracts of chromatograms obtained from non-chiral GC-FID analyses of in vitro reactions (conditions summarized in **Table S5**) are represented in **Figures S3** and **S4**, below.

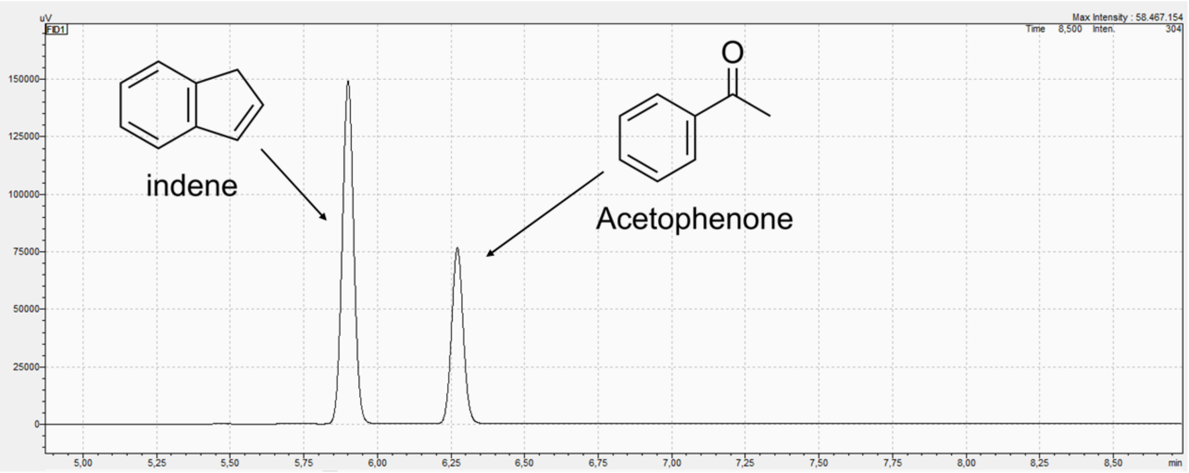

**Figure S3.** Extract from chromatogram obtained from GC-FID analysis with parameters from **Table S5**. The peaks representing indene and acetophenone are indicated. RT: 5.90 min indene, 6.27 min acetophenone.

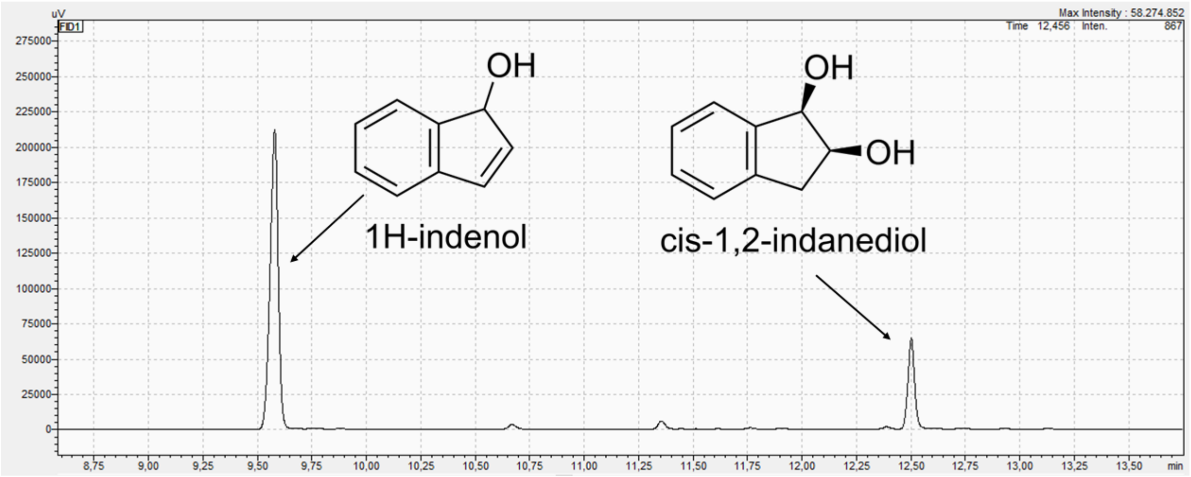

**Figure S4.** Extract from chromatogram obtained from GC-FID analysis with parameters from Table S5. The peaks representing the products 1*H*-indenol and *cis*-1,2-indanediol are indicated. RT: 9.58 min 1*H*-indenol, 12.50 min *cis*-1,2-indanediol.

#### Quantification of $H_2O_2$ via ABTS-HRP assay

The quantification of $H_2O_2$ in aqueous reaction solutions was performed via an ABTS-HRP assay. Therefore, a calibration curve was established with defined concentrations of $H_2O_2$. The calibration curve obtained by plotting the concentration of $H_2O_2$ (0-500 µM) against the absorbance measured at 414 nm is shown in **Figure S5**.

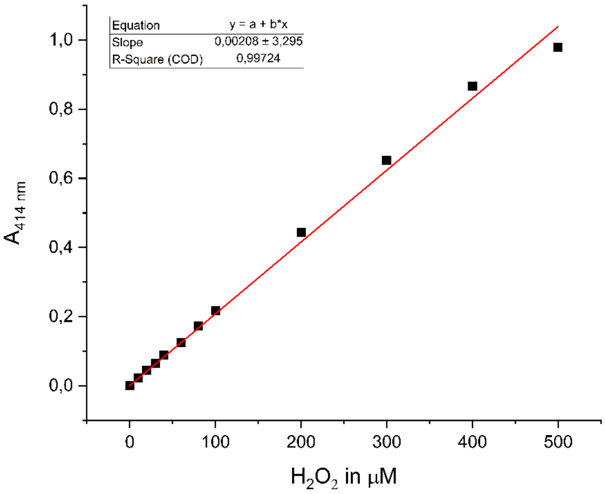

**Figure S5.** Calibration curve for $H_2O_2$ determination via ABTS-HRP assay measuring the absorption at 424 nm. The linear regression is indicated.

### Time-Course Experiments

Product formation over time was evaluated for the CDO-WT and CDO-PDR Hybrid system using GC-FID (GC parameters can be found in **Table S5**). Determined product formations plotted against time are depicted in **Figure S6**. As indicated in the figures, a activity of 94.47 mU for CDO-PDR (**Figure S6a**) and 35.33 mU (**Figure S6b**).

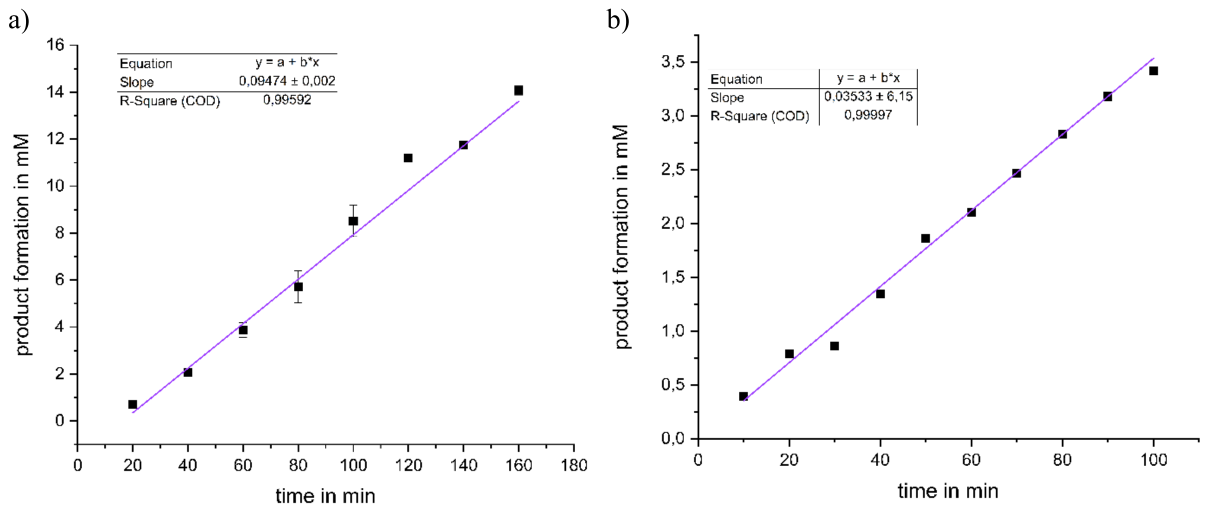

**Figure S6.** Product formation over time is determined in *in vitro* reactions using **a)** CDO-PDR hybrid system or **b)** CDO-WT, respectively. Product formation corresponds to the total amount of **1a** and **1b**. Linear equations are indicated in the figures.
